# Pulsed broad-spectrum UV light effectively inactivates SARS-CoV-2 on multiple surfaces

**DOI:** 10.1101/2021.02.12.431032

**Authors:** Alexander S. Jureka, Caroline G. Williams, Christopher F. Basler

## Abstract

The ongoing SARS-CoV-2 pandemic has resulted in an increased need for technologies capable of efficiently disinfecting public spaces as well as personal protective equipment. UV light disinfection is a well-established method for inactivating respiratory viruses. Here, we have determined that broad-spectrum, pulsed UV light is effective at inactivating SARS-CoV-2 on multiple surfaces. For hard, non-porous surfaces we observed that SARS-CoV-2 was inactivated to undetectable levels on plastic and glass with a UV dose of 34.9 mJ/cm^2^ and stainless steel with a dose of 52.5 mJ/cm^2^. We also observed that broad-spectrum, pulsed UV light is effective at reducing SARS-CoV-2 on N95 respirator material to undetectable levels with a dose of 103 mJ/cm^2^. We included UV dosimeter cards that provide a colorimetric readout of UV dose and demonstrated their utility as a means to confirm desired levels of exposure were reached. Together, the results present here demonstrate that broad-spectrum, pulsed UV light is an effective technology for the inactivation of SARS-CoV-2 on multiple surfaces.

## Introduction

In late 2019, the novel severe acute respiratory distress syndrome virus 2 (SARS-CoV-2) emerged from Wuhan, China [1,2]. SARS-CoV-2 is a member of the *Coronaviridae* family of enveloped negative-sense RNA viruses. It is classified in the *Betacoronavirus* genus of which other notable members are the highly pathogenic SARS-CoV and the Middle East respiratory syndrome virus (MERS-CoV) [3]. Since its emergence, SARS-CoV-2 has been the cause of the most severe pandemic in the last century. Despite significant efforts to contain the spread of SARS-CoV-2, as of February 7^th^, 2021 it has caused over 105 million cases and resulted in over 2.3 million deaths worldwide [4].

High case counts raise concerns about infections arising from contaminated public spaces, such as mass transit vehicles and hospital spaces that have housed SARS-CoV-2 positive patients. The need for effective means to eliminate SARS-CoV-2 from environmental surfaces is supported by studies demonstrating the capacity of the virus to survive on a variety of surfaces for significant periods of time on a variety of surfaces. For example, infectious virus could be recovered from plastic or steel for 72 hours and on cardboard after 24 hours [5]. There is also substantial evidence that SARS-CoV-2-infected individuals shed virus into their environment. Analysis by RT-PCR of COVID-19 patient rooms and other hospital settings demonstrated frequent contaminating viral RNA on surfaces [6-9]. In some studies, however, lower frequency of surface contamination has been reported [10]. Outside of the healthcare setting, rooms of cruise ship passengers who had COVID-19 were also contaminated with viral RNA [11]. Viral RNA has also been found on various surfaces in households with SARS-CoV-2 infected individuals [12,13].

Given the potential for fomite transmission, the World Health Organization has provided guidance on cleaning and disinfection where SARS-CoV-2 contamination could occur [14]. Because of the importance of respiratory protection and shortages of personal protective equipment, methods to disinfect and reuse N95 filtering respirators has also been of significant interest [15,16]. UV light as well as other methods have been either proposed or tested as a means to disinfect N95 masks and other PPE [16-24].

While chemical disinfectants and alcohols are effective methods of inactivating SARS-CoV-2 in most circumstances, disinfection of large spaces using these methods is a laborious process requiring close contact with potentially contaminated surfaces [25-28]. UV light has long been established as an effective and direct method for the inactivation of enveloped viruses [29]. UV disinfection approaches provide a significant advantage as they are less laborious to employ and do not necessarily require close contact with potentially contaminated surfaces.

Here, we report the efficacy of broad-spectrum, pulsed UV light in inactivating SARS-CoV-2 on glass, plastic, stainless steel, and N95 respirator material. Additionally, we have tested the effectiveness of UV dosimeter cards that would provide end users the ability to quickly determine if a high enough dosage of UV light has been applied to a surface. Together, the data reported here demonstrate that broad-spectrum, pulsed UV light is highly effective at inactivating SARS-CoV-2 on multiple surfaces.

## Materials and Methods

### Cells and virus

Vero E6 cells (ATCC# CRL-1586) were maintained in DMEM supplemented with heat-inactivated fetal bovine serum (FBS; Gibco). SARS-CoV-2, isolate USA_WA1/2020, was obtained from the World Reference Collection for Emerging Viruses and Arboviruses at the University of Texas Medical Branch. SARS-CoV-2 virus stocks were propagated as previously described [30].

### Surface Inoculation

The following surfaces were tested within the wells of a 24 well plate in triplicate: glass coverslips, 0.5×0.5 cm stainless steel squares, the tissue culture plate well (plastic; polystyrene), and 0.5×0.5 cm squares of N95 respirator material from a 3M™ 9210+ respirator. Four 0.5×0.5 cm squares of UV dosimeter cards (Intelligo Technologies) were placed in each corner of the plate to confirm even UV exposure across the plates and allow comparison of the cards to a UV dosage meter as a means to quantify exposure dose (Supplemental Figure 1). For surface inoculation, 12 µL of SARS-CoV-2 stock virus (USA_WA1/2020; 8.3×10^4^ pfu) in OptiMEM supplemented with 1x antibiotic/antimycotic (Gibco) was pipetted directly onto each surface being tested and spread with a pipette tip to facilitate efficient drying. Surface samples inoculated with virus were allowed to dry in the 24 well plates with the lids off for 1 hour at room temperature in the biosafety cabinet. After the surfaces had dried, the 24-well plate lids were replaced, and all plates not being exposed to UV were placed inside a black opaque container in a separate biosafety cabinet to avoid incidental UV exposure.

**Figure 1.**
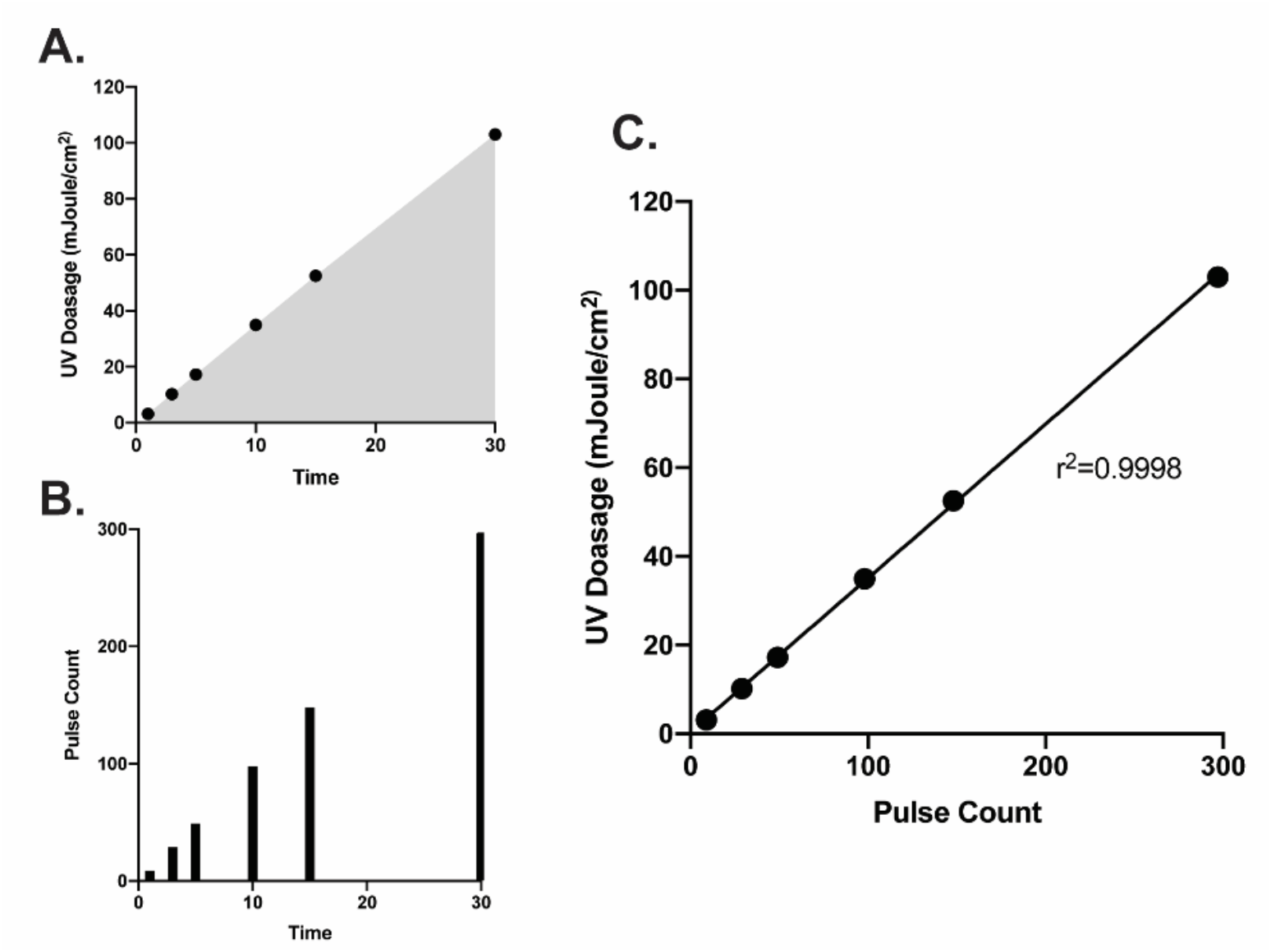
UV dosage (A) and pulse-counts (B) recorded over time from the Helo F2 device. (C) Pulse count plotted versus UV dosage shows a significant positive correlation. Data were fit with a linear regression in GraphPad prism.

### UV exposures

The Puro UV Helo F2 device was placed in the center of the biosafety cabinet. The test samples as well as an UV dosage meter (ILT2500; International Light Technologies) equipped with a calibrated SED270 detector were placed 1 meter away facing the Helo F2 device. Care was taken to ensure that the UV dosage meter and surfaces being exposed were as inline as possible before beginning testing. The 24-well plates containing the test surfaces were positioned so that the plates were nearly vertical (∼85 degrees) and approximately 3 inches above the surface of the biosafety cabinet to avoid shadowing (Supplemental Figure 2). Once the 24-well plate was positioned, the UV dosage meter was zeroed to account for ambient UV. Once zeroed, the UV dosage meter was set to “integrate” mode to measure UV dosage over time and total pulse-counts. The Helo F2 device was initiated using an electronic timer set to the indicated exposure time with an additional minute added to account for device startup procedures. UV dosage for a given timepoint was recorded as mJ/cm^2^ along with the corresponding pulse-count. UV dosimeter cards were collected and photographed to record the color change. One card from the 3-minute timepoint was lost due to airflow in the biosafety cabinet. For the purposes of graphical depiction, UV dosimeter card color post-UV exposure was replicated using the eyedropper tool in Adobe Illustrator.

**Figure 2.**
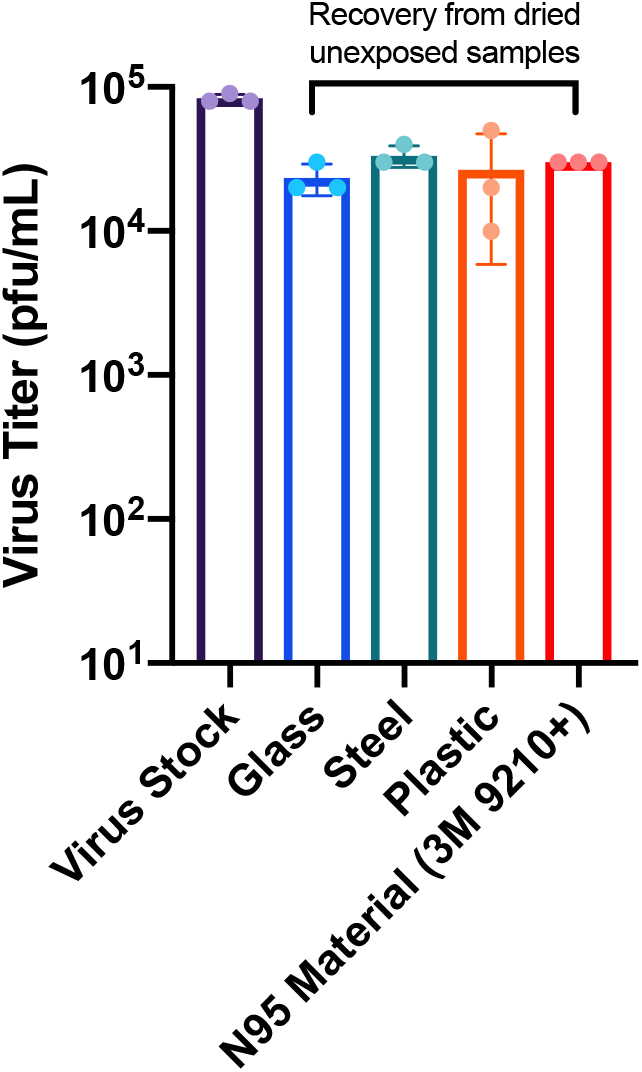
Virus titer reduction due to drying. Virus stock represents the amount of virus initially inoculated onto the surfaces. Glass, steel, plastic, and 3M N95 material samples represent the virus titer recovered from dried, unexposed samples after harvesting.

### Sample harvesting

After all exposures were complete, 1 mL of sterile PBS was placed inside the surface-containing wells and allowed to rehydrate for 15 minutes before transferring to a 1.5 mL centrifuge tube and storing at −80C for further analysis. Additionally, 12 µL of stock virus (USA_WA1/2020; 8.3×10^4^ pfu) was added to 1 mL of sterile PBS and stored at −80C at the same time as the other samples in order to control for the loss of virus titer due to the drying process.

### Virus quantification by plaque assay

Vero E6 cells were plated to confluency in 24-well plates 24 hours prior to infection. Ten-fold serial dilutions of SARS-CoV-2 containing samples were added onto the cells (100 µL) and virus was adsorbed for 1 hour with shaking at 15-minute intervals. After the adsorption period, 1 mL of 0.6% microcrystalline cellulose in DMEM supplemented with 2% fetal bovine serum and 1x antibiotic-antimycotic was overlaid onto to the cells and plates were incubated at 37C/5% CO2 for 72 hours, as described previously [30]. After incubation, the microcrystalline cellulose overlay was aspirated from the well, and cells were fixed with 10% neutral buffered formalin for 1 hour at room temperature. Plates were then washed with water and stained with crystal violet to visualize plaques. Plaques were quantified and recorded as plaque forming units per mL (pfu/mL). All samples assayed were only subjected to one freeze-thaw cycle.

## Results

### UV dosage and pulse counts show significant correlation

We first determined the range of UV dosage that could be achieve with the Helo F2 device between 1 and 30 minutes of exposure. The device produces a theoretical 10 pulses per minute. In practice, we achieved 6 pulses after 1 minute and 297 pulses after 30 minutes, corresponding to cumulative UV doses of 3.14 mJ/cm^2^ and 103 mJ/cm^2^, respectively (Figure 1A, B). Interestingly, we observed the UV dosage output by the Helo F2 device over time has a significant and linear correlation with the overall pulse count (Figure 1C). This suggests that it is possible to identify a specific amount of time required for inactivation depending on the dosage required. Additionally, the exposure times and UV dosage range tested in this study encompass previously reported effective UV exposure times and doses for inactivating SARS-CoV-2 with similar UV devices [31,32].

### Broad spectrum pulsed UV light effectively inactivates SARS-CoV-2 on multiple surfaces

To determine the effectiveness of the Helo F2 device in inactivating SARS-CoV-2, glass, stainless steel, plastic, and N95 respirator material inoculated with SARS-CoV-2 (see materials and methods) were exposed to pulsed UV light for 1, 3, 5, 10, 15, and 30 minutes from a distance of 1 meter. Time zero represents samples that were inoculated, dried and quantified for infectivity without UV exposure. SARS-CoV-2 titers recovered from the unexposed UV controls indicate that the drying process resulted in an approximately 3-fold decrease in titers when compared to the same inoculums that had not been dried prior to titration (Figure 2). Additionally, similar amounts of SARS-CoV-2 were recovered from all surfaces, indicating that the recovery process was efficient for all tested surfaces.

For the UV exposed samples, we observed that for the hard, non-porous surfaces (glass, stainless steel, and plastic) a pulse-on time of 5 minutes (17.2 mJ/cm^2^) was sufficient to achieve a 3-log_10_ reduction of infectious virus recovered from the surfaces (Figure 3A-C). Exposure for 10 minutes (34.9 mJ/cm^2^) was sufficient to reduce infectious virus to nearly undetectable levels on glass and plastic, while 15 minutes (52.5 mJ/cm^2^) was required for the same effect on stainless steel. N95 respirator material (3M 9210+) required 15 minutes of exposure to achieve a 2.86 log10 reduction in infectious virus and 30 minutes (103 mJ/cm^2^) exposure for reduction to undetectable levels which likely due to the porous and multilayer structure of N95 material (Figure 3D) [33-35]. Taken together, these data suggest that broad-spectrum pulsed UV light is capable of effectively inactivating SARS-CoV-2 in short periods of time on hard non-porous surfaces, and on N95 respirator material with longer exposure times.

**Figure 3.**
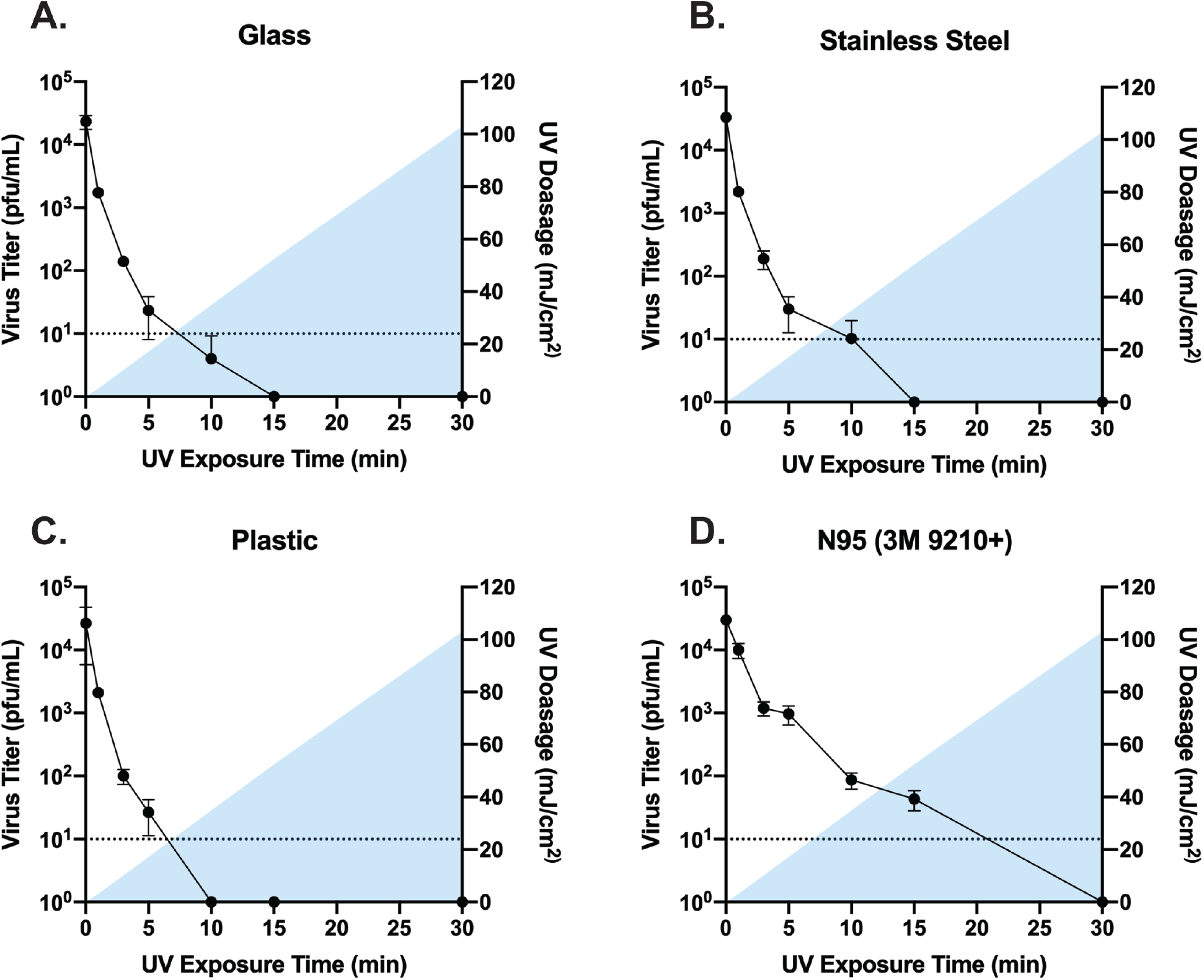
Titers of infectious SARS-CoV-2 recovered from UV exposed glass, stainless steel, plastic, and N95 material (A-D). Time 0 represents controls that were not exposed to UV. All timepoints are representative of the mean and standard error of 3 replicates. Blue shading represents the area under the curve for the UV dosage acquired over time. Samples with data points below the limit of detection resulted from a subset of datapoints having undetectable levels of virus. Undetected samples were assigned a value of 1 for graphing purposes.

**Figure 5.**
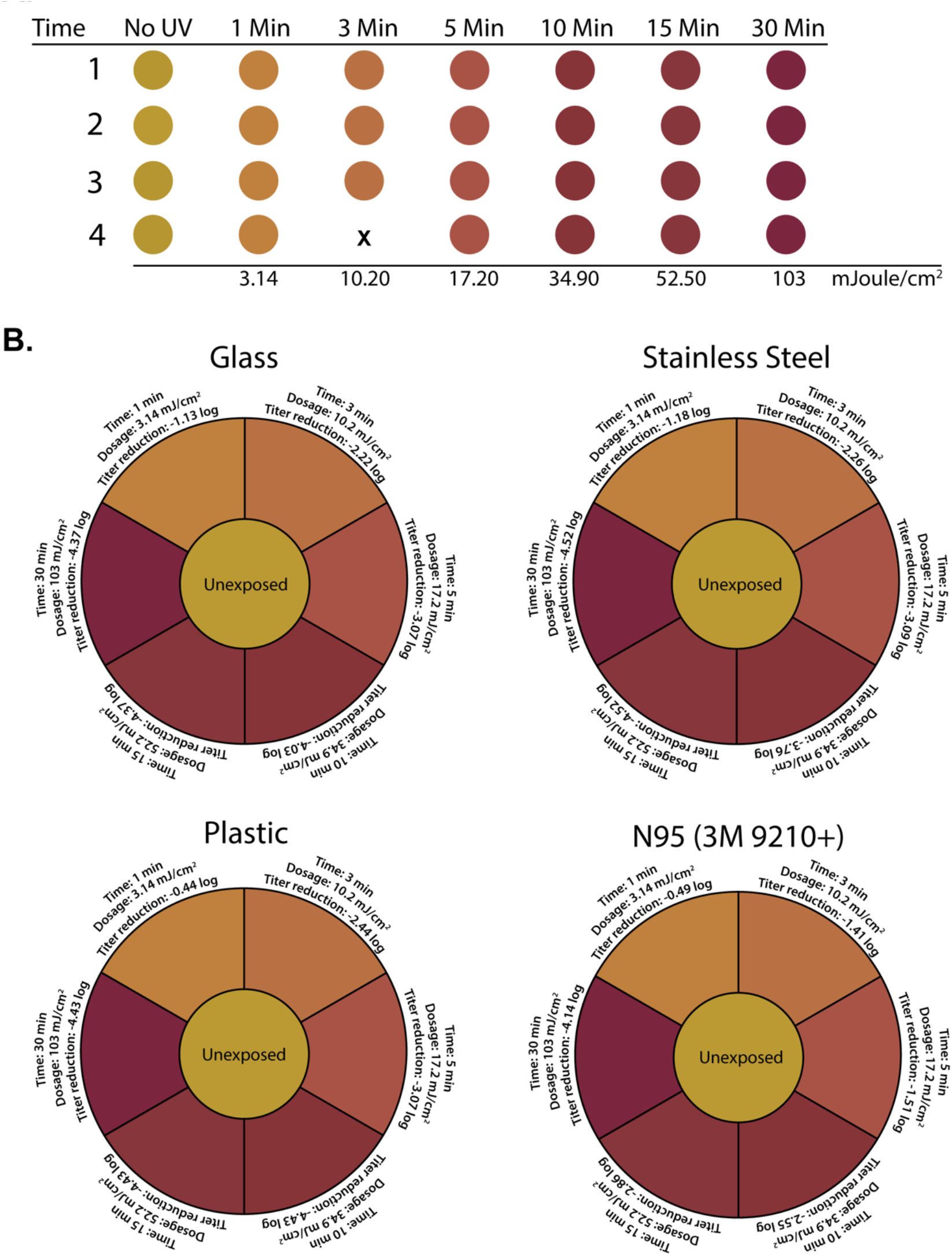
(A) Colorimetric change of indicator cards at indicated timepoints and dosages. (B) Compilation of time, dosage, and SARS-CoV-2 titer reduction data as a function of indicator card color. Indicator color in A and B is graphically depicted by using the eyedropper color tool within Adobe illustrator on recovered UV indicator cards.

### Correlating colorimetric UV dosimeter cards to physical UV dosimeter and virus titer reductions

From a point-of-use perspective, electronic UV dosimeters like the one employed here are expensive and require manual set-up and operation, making them a less than ideal option for ensuring the correct dosage has been applied to a surface. Because of this, UV dosimeter cards are available that exhibit a colorimetric change in response to increasing doses of UV light. In tandem with testing the effectiveness of broad-spectrum UV light in inactivating SARS-CoV-2, we were also interested in determining the functionality of UV dosage cards specifically designed for pulsed-UV light sources. To test their functionality, 4 test pieces of experimental UV dosage card material were included in each UV exposure timepoint. We determined that these functioned as intended with a significant color change occurring with increasing doses of UV (Figure 4A). We identified that the color change was even across all cards recovered at each timepoint indicating a high degree of reproducibility across the material (Supplemental Figure 3). Given that these cards would be intended for end-users to ensure that a high enough dosage had been applied for inactivation, we compiled our UV dosage meter and SARS-CoV-2 titer reduction data to correlate with the color change from the cards (Figure 4B). Taken together this UV reactive card material represents an effective alternative to high-cost dosage meters for end users of broad-spectrum pulsed UV disinfection equipment.

## Discussion

The availability of information regarding the inactivation of SARS-CoV-2 for environmental disinfection is of paramount importance. UV disinfection is a validated technology that has been utilized for decades to inactivate pathogens on surfaces, as well as in air and water [36,37]. The effectiveness of UV light, pulsed or constant, in inactivating pathogens is expected to be directly related to the dosage applied [33]. Here, we have described empirically determined dosages of broad-spectrum, pulsed UV light that are effective for the inactivation of SARS-CoV-2 on multiple relevant surfaces. Additionally, we have also demonstrated the effectiveness of colorimetric UV dosimeter cards for dosage determination by the end-user.

While only a single device was tested in this study, a variety of UV inactivation studies have been published focusing on SARS-CoV-2 and other coronaviruses using different wavelengths and environmental conditions [38-43]. This has led to the development of new UV disinfection products in an effort to address proper disinfection of PPE and surfaces as the pandemic continues [44]. Disinfection of potentially contaminated surfaces by broad-spectrum UV light is a particularly attractive option when compared to chemical disinfectants as its less laborious and does not require close contact with contaminated surfaces. Additionally, with PPE shortages being an ongoing concern, our data demonstrates that broad-spectrum pulsed UV could be an effective strategy for the disinfection of N95 respirators, although our study did not determine how pulsed UV exposure affects N95 performance.

While our data demonstrate that broad-spectrum pulsed UV light is an effective method for inactivating SARS-CoV-2, one notable limitation of our study is that surfaces were only exposed to the UV light at a distance of one meter. However, our data demonstrated that colorimetric UV dosimeter cards work as expected and provide a clear indication of the dosage being applied to a particular surface where the card is in place. In the event that a surface is more than one meter from a given UV device, using our data as a reference the inverse square law could be applied to determine the amount of time required to achieve a particular dosage at the necessary distance [45-47]. In addition, utilizing UV dosimeter cards like those tested here would provide a rapid, low-cost method for testing broad-spectrum pulsed UV light devices at different distances to ensure that an effective dosage is delivered.

The data presented here demonstrate that broad-spectrum UV light is an effective means of inactivating SARS-CoV-2 on multiple surfaces, including N95 respirator material. Additionally, UV dosimeter cards like those tested here represent an effective and straightforward means for point-of-care users of UV disinfection equipment to ensure that surfaces have been properly disinfected.

## Acknowledgments and Competing Interests

We would like to acknowledge the GSU High Containment Core for their assistance in performing these studies. PURO UV Disinfection Lighting provided the Puro UV Helo F2 device, the UV dosimeter cards, the UV meter and funds for the studies described. All data and analyses were collected and performed without input from PURO UV Disinfection Lighting.

