## Supplemental Figure 1, Supplemental Figure 2, Supplemental Figure 3 for "Pulsed broad-spectrum UV light effectively inactivates SARS-CoV-2 on multiple surfaces"

**Supplemental Figures**

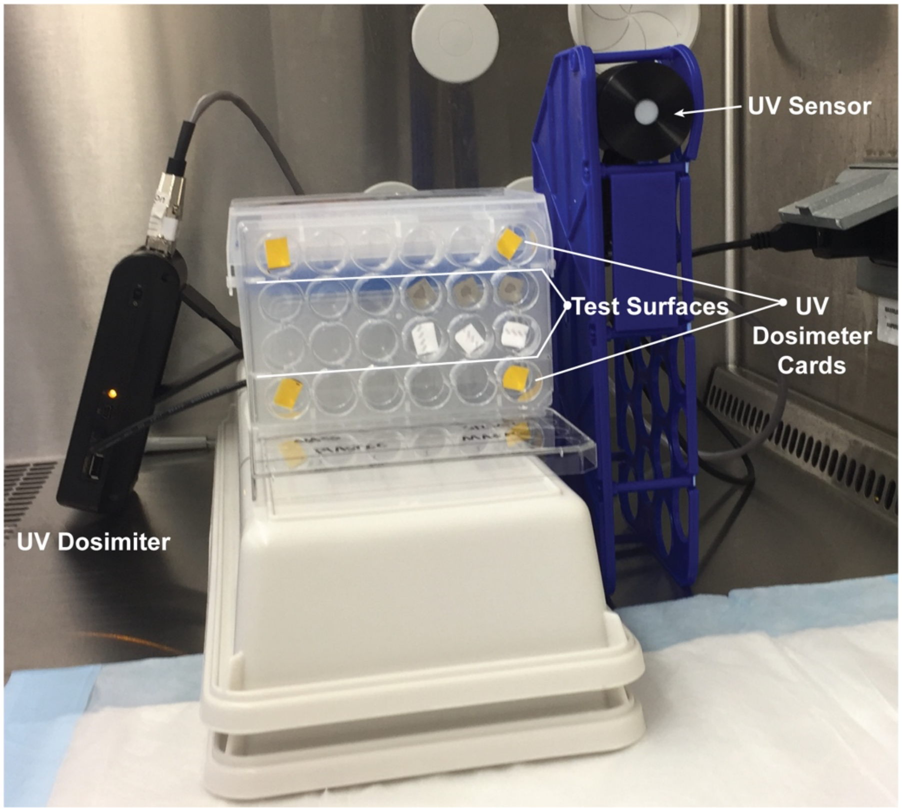


**Supplemental Figure 1.** Test plate arrangement.

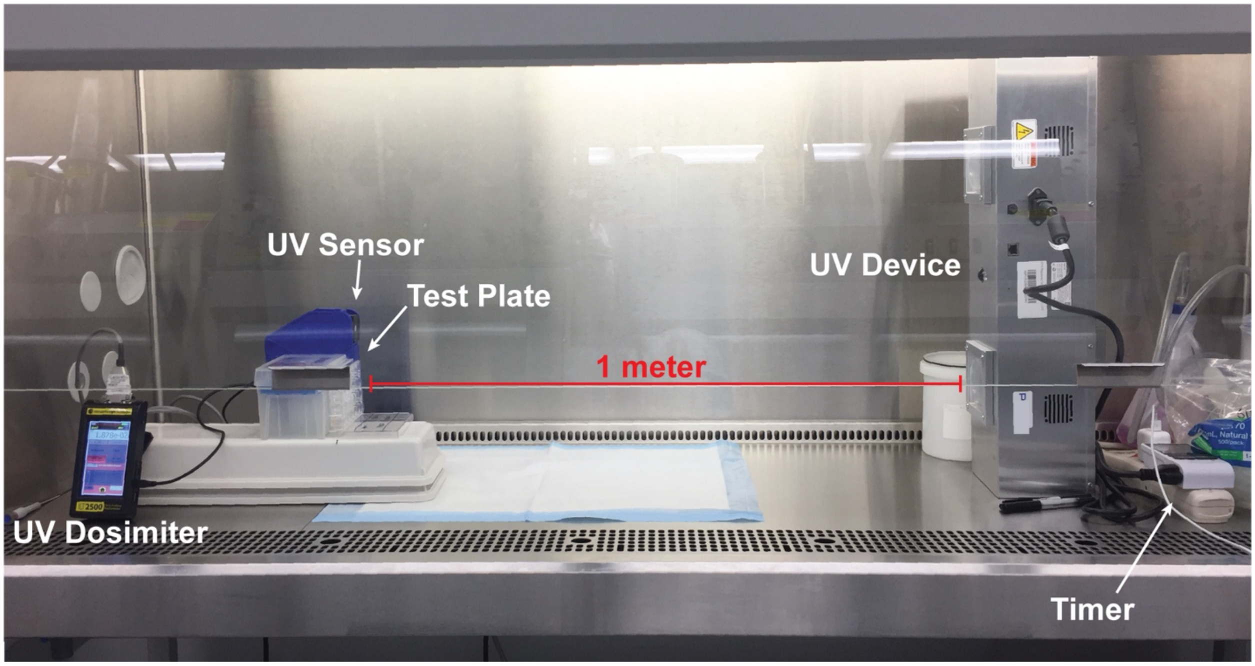


**Supplemental Figure 2.** Experimental setup.

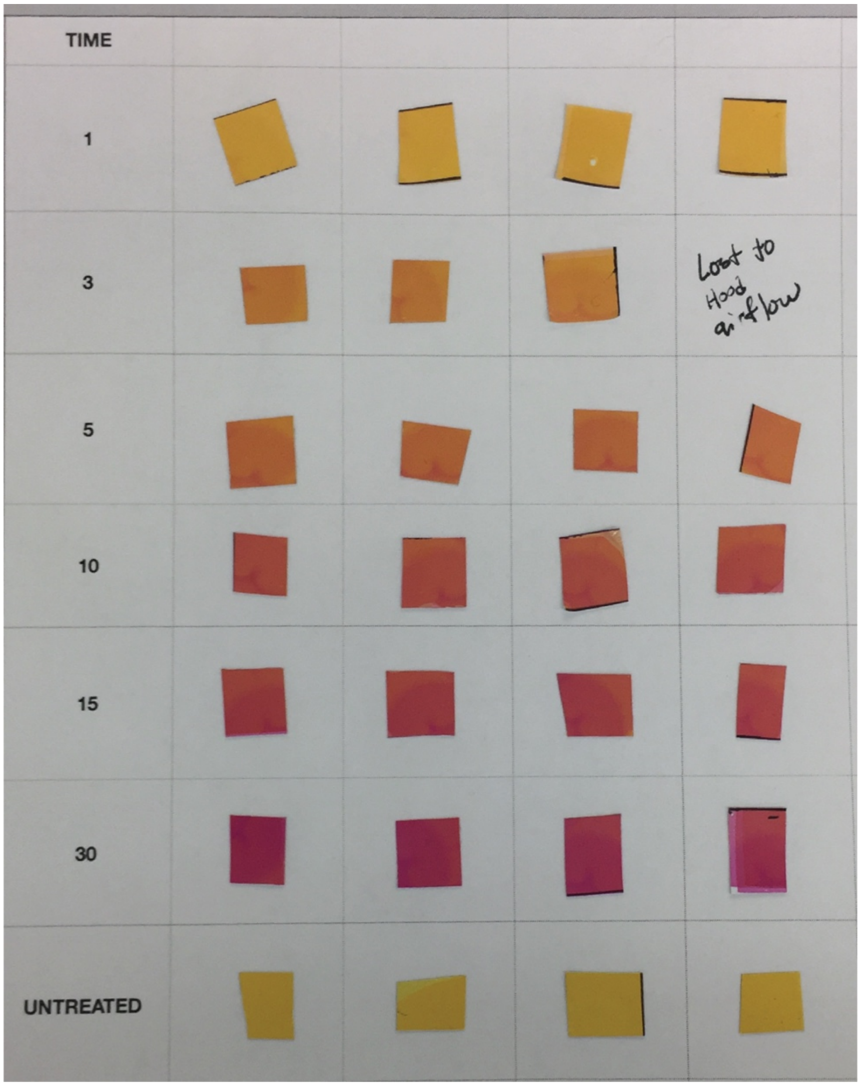


**Supplemental Figure 3.** UV dosimeter cards collected from each test plate.
